# Development of the Brain Functional Connectome Follows Puberty-Dependent Nonlinear Trajectories

**DOI:** 10.1101/2020.09.26.314559

**Authors:** Zeus Gracia-Tabuenca, Martha Beatriz Moreno, Fernando Barrios, Sarael Alcauter

## Abstract

Adolescence is a developmental period that dramatically impacts body and behavior, with pubertal hormones playing an important role not only in the morphological changes in the body but also in brain structure and function. Understanding brain development during adolescence has become a priority in neuroscience because it coincides with the onset of many psychiatric and behavioral disorders. However, little is known about how puberty influences the brain functional connectome. In this study, taking a longitudinal human sample of typically developing children and adolescents (of both sexes), we demonstrate that the development of the brain functional connectome better fits pubertal status than chronological age. In particular, centrality, segregation, efficiency, and integration of the brain functional connectome increase after the onset of the pubertal markers. We found that these effects are stronger in attention and task control networks. Lastly, after controlling for this effect, we showed that functional connectivity between these networks is related to better performance in cognitive flexibility. This study points out the importance of considering longitudinal nonlinear trends when exploring developmental trajectories, and emphasizes the impact of puberty on the functional organization of the brain in adolescence.

**Significance Statement:** Understanding the brain organization along development is a crucial challenge for Neuroscience. In particular, during adolescence there is a great impact in body and cognitive functions as well as substantial incidence of mental health disruptions. Here, we tested how brain organization changes along this period based on the properties of the functional connectome in a longitudinal pediatric sample. We found a nonlinear increase in the connectivity and the brain network efficiency, particularly after the onset of puberty. These effects were more prominent in association networks. In addition, higher connectivity in those areas was associated with better performance in cognitive flexibility. These results demonstrate the importance of considering pubertal assessment as well as nonlinear trends in developmental studies.

## 1. Introduction

Adolescence is a developmental period that dramatically impacts body and behavior. Pubertal hormones play an important role in adrenal, gonadal, and growth axes, having a substantial impact not only in the morphological changes in the body but also in brain structure and function (Vijayakumar et al., 2018). Therefore, measuring brain development along this stage is complicated given that these changes are not only influenced by chronological age (Blakemore et al., 2010). Other factors play a role, such as sex, ethnic, and environmental influences, and all of them with a high degree of variability between individuals (Sawyer et al., 2018). Also, adolescence is considered a critical period for mental health, when many behavioral and psychiatric disorders first manifest (Kessler et al., 2005; Paus et al., 2008). Additionally, cognitive abilities as important as task-switching or cognitive flexibility strengthen in adolescence (Somsen, 2007; Hauser et al., 2015), together with other executive functions, including working memory, inhibition, and attention (Gur et al., 2012). Such cognitive improvement, that usually starts in late infancy and continues improving until early adulthood, may result from the morphological and metabolic changes of the brain, which are expected to improve their efficiency to solve complex problems (Baum et al., 2017; L. R. Chai et al., 2017).

The widespread use of in vivo neuroimaging techniques has shifted the study of developmental trajectories from focal brain areas to a system perspective (Di Martino et al., 2014; Zuo et al., 2017). Thus, modeling the brain as a set of interconnected elements, or connectome, allows the description of its functional organization in terms of network-based properties, such as the functional segregation and integration, and the efficiency of the brain network (Rubinov and Sporns, 2010; Stam and Van Straaten, 2012). This framework has been extensively applied in other lifespan studies, showing dramatic developmental changes in relation with chronological age (Zuo et al. 2011; Tomasi and Volkow, 2012; Alcauter et al., 2014; Betzel et al., 2014; Gao et al., 2015; Gracia-Tabuenca et al., 2018). Particularly in adolescence, previous cross-sectional studies have shown an increase in functional segregation along with cognitive specialization (Fair et al., 2009; Satterthwaite et al., 2013a; S. Gu et al., 2015), whereas other studies have emphasized an increase in the functional integration between brain networks with age (Hwang et al., 2012; Marek et al, 2015). Such potentially divergent results may be influenced by differences in samples and/or methodologies; however, the implementation of longitudinal samples and/or nonlinear trends will potentially reduce the bias when characterizing neurodevelopmental trajectories (Zuo et al., 2017; Vijayakumar et al., 2018). Additionally, the motion artifact is another potential bias given that younger subjects tend to move more in pediatric populations affecting the sample inferences (Satterthwaite et al., 2013a). Previous findings have described mitigating strategies at the edgewise level (Ciric et al., 2017), however little is known regarding functional organization measures in a developmental context.

Furthermore, the pubertal spurt is an important source of variability in adolescence when considering developmental trajectories based on chronological age, but it is rarely considered (Blakemore et al., 2010; Vijayakumar et al., 2018). A pair of recent studies have taken an important step in this direction, by fitting the developmental trajectories of fronto-subcortical connectivity with the pubertal stage (Spielberg et al., 2014; van Duijvenvoorde et al., 2019). However, the relationship between the pubertal stage and the developmental of the brain functional organization still remains largely unexplored.

This work is focused on elucidating the aforementioned discrepancies and unknowns. First, we tested how mitigation strategies of the motion artifact may impact the functional connectome features. Then, we tested the development of the functional organization of the adolescent brain, by exploring the longitudinal trajectories with generalized additive mixed models, and their relation with pubertal stage and cognitive development.

## 2. Methods

### 2.1 Sample

The total sample consisted of 98 typically developing subjects (45 male, ages from 6.71 to 18.06 years old), of whom 41 and 16 returned to a second and a third session, respectively (Figure 1). A general invitation describing the study protocol and inclusion criteria was sent to local schools. Inclusion criteria included full-term gestation (at least 37 weeks) and enrollment in its corresponding academic year. Exclusion criteria consisted of any neurological or psychiatric disorder identified with a semi-structured interview: MINI-KID for minors and MINI for those who were older than 18 years old. Written informed consent was required, and in the case of minors, parents were required to sign the consent and children to verbally assent. Study protocols followed the Declaration of Helsinki and were approved by the Institutional Review Board. Follow-ups occurred after 5 years and the second after 2 years.

**Figure 1.**
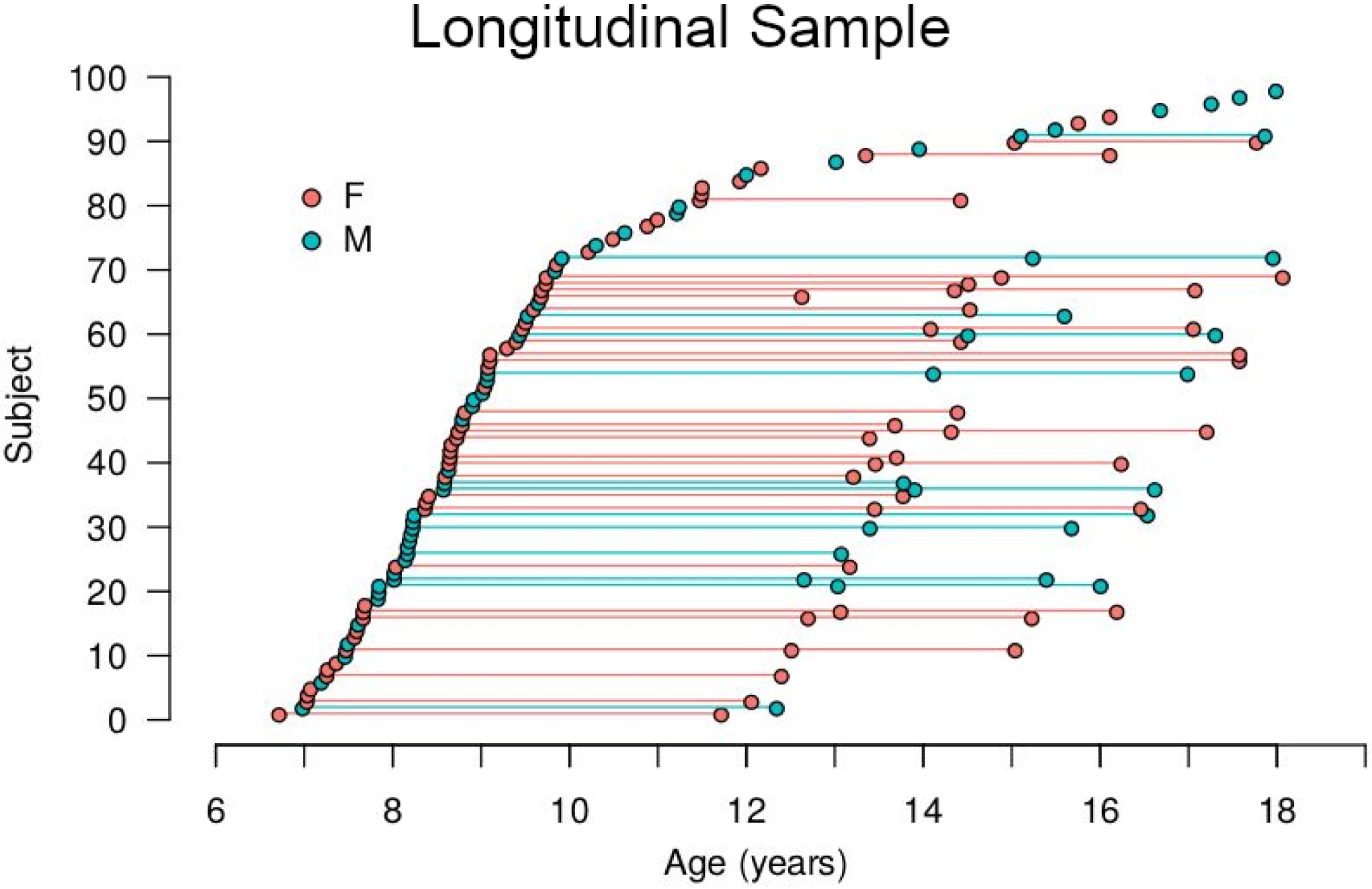
Longitudinal sample. Each dot represents a subject at the age of assessment. Lines represent longitudinal assessments of a subject. Abbreviations: female (F), male (M).

### 2.2 Pubertal status

Participants completed the Pubertal Development Scale (PDS; Petersen et al., 1988). PDS is a self-report questionnaire that encompasses Likert-like scales to respond about growth spurt in height, pubic hair, and skin change for both sexes; plus facial hair growth and voice change for males, and breast growth and menarche for females, resulting in a four-level outcome: 1) absence, 2) first signs, 3) evident, and 4) finished pubertal spurt. This scale allows taking into account sex-specific patterns of puberty development even at the same chronological age. Children under 10 years old were set to level 1 of the PDS, as previous studies have shown similar values at these ages (Hibberd et al., 2015; van Duijvenvoorde et al., 2019). Missing values corresponding to 8 time-points (4 females) were estimated with two age-smooth splines by sex with a Generalized Additive Mixed Model (GAMM) following van Duijvenvoorde et al. (2019).

### 2.3 MRI sessions

MRI sessions consisted of a resting-state fMRI sequence followed with a high-resolution T1-weighted scan. 150 whole-brain fMRI volumes were acquired using a gradient recalled T2* echo-planar imaging sequence (TR/TE = 2000/40 ms, voxel size 4 × 4 × 4 mm^3^). Participants were instructed to keep their eyes closed and not to fall asleep. To facilitate participants to remain awake, this sequence was always applied in the morning and at the beginning of the MRI session. Furthermore, participants were asked to avoid staying up after their usual bedtime. Structural T1 sequence was obtained using a 3D spoiled gradient recalled (SPGR) acquisition (TR/TE = 8.1/3.2 ms, flip angle = 12.0, voxel size 1×1×1 mm^3^). MR images were acquired with a 3T MR GE750 Discovery scanner (General Electric, Waukesha, WI), using an 8-channel-array head coil. 20 sessions were acquired with a 32-channel coil, therefore a dummy covariate was included in the subsequent analyses.

### 2.4 Preprocessing

Anatomical T1 images were denoised with non-local means (Manjón et al., 2010) and N4 bias field correction (Tustison et al., 2010). fMRI preprocessing was implemented using FMRIB’s Software Libraries (FSL v.5.0.6; Jenkinson et al., 2012; RRID: SCR_002823) including slice timing, head motion correction, brain extraction, intensity normalization, regression of confounding variables, spatial normalization, and band-pass temporal filtering (0.01–0.08 Hz). Each fMRI dataset was registered to its corresponding structural image with a rigid-body transformation, followed by a two-step diffeomorphic SyN registration (Avants et al., 2008; RRID: SCR_004757) to a pediatric brain atlas (NIHPD 4.5-18.5; Fonov et al., 2011), and then to the MNI-152 standard template.

Framewise displacement was estimated and summarized by the means of FD-RMS computed with FSL’s MCFLIRT (Jenkinson et al., 2002), and those volumes with more than 0.25 mm were considered “spikes”, and they were regressed out from the fMRI time series. Given that pediatric samples tend to move more inside the scanner than adult populations, we implemented two rigorous confounding regression strategies, one including the global signal regression (GSR) and the other one without GSR:

- GSR: 36 parameters were regressed out (Satterthwaite et al., 2013a), including the time series, derivatives, and their corresponding quadratic terms of the six estimated motion parameters, and the average signal from global and the eroded white matter (WM) and cerebrospinal fluid (CSF).
- no-GSR: 37 parameters were regressed out (Gracia-Tabuenca et al., 2020), including the time series, derivatives, and their corresponding quadratic terms of the six estimated motion parameters, and the average signal from the eroded WM and CSF, plus five principal components of the WM+CSF, this last procedure is termed *aCompCor* (X. J. Chai et al., 2012; Muschelli et al., 2014).

Eighteen sessions with less than four minutes of “spikes”-clean time series were discarded for further analyses (Satterthwaite et al., 2013a), resulting in a final sample of 89 typically developing subjects (39 male, age: 6.7 – 18.1 years old), of which 37 and 11 returned for a second and third session, respectively.

### 2.5 Network features

Brain network construction was based on the 264 regions of interest (ROIs) described by Power et al. (2011), which consist of 5-mm radius spheres, encompassed in 13 different functional networks. This atlas has been widely applied in pediatric studies (Satterthwaite et al., 2013a, 2014; S. Gu et al., 2015; Marek et al., 2015; Ciric et al., 2017; L.R. Chai et al., 2017; Gracia-Tabuenca et al., 2020). Edges were calculated as the functional connectivity between every pair of ROIs, via the Pearson’s cross-correlation from their average time series. Based on these matrices, weighted graph theory metrics were computed to characterize network organization features. The sum of edges or weighted degree (*d*), clustering coefficient (*C*), efficiency (*E*), and characteristic path length (*L*) were estimated to account for centrality, segregation, and integration in terms of functional network organization (Rubinov & Sporns, 2010). *C* was calculated based on the Barrat formula (Barrat et al., 2004). *L* was calculated as the average shortest-paths based on the Dijkstra algorithm (Dijkstra, 1959), where pairwise distances were set as the inverse of the corresponding functional connectivity edge. *E* was computed as the average of the inverse of the shortest-paths of the connectome. For *L*, the shortest-path of disconnected nodes was set to the maximum observed value of the network in order to avoid infinite outcomes (Fornito et al., 2010). Barrat and Dijkstra’s algorithms were calculated via the R package ‘*igraph*’ (Csardi & Nepusz, 2006; https://CRAN.R-project.org/package=igraph).

Additionally, we tested the relationship of the head motion (i.e., FD-RMS) and the brain network properties via linear mixed-effects (LME) models for the two confounding regression strategies, in order to identify the best preprocessing strategy, as suggested by Ciric et al. (2017). Besides, we tested if such relationships are affected by the Euclidean distance of the network’s nodes (Satterthwaite et al., 2013b).

### 2.6 Neurodevelopmental trajectories

Generalized Additive Mixed Models (GAMM) were applied to account for the non-linear trends (smooth splines) and longitudinal effects, including head-coil and average head-motion as covariates. Four different models were tested: age, PDS, age-by-sex, and PDS-by-sex. Basis functions of the age or PDS splines were set to a maximum of four (k = 4) according to van Duijvenvoorde et al. (2019). Mixed-effects were estimated via Maximum Log-likelihood in order to compare between models. The best model was selected according to the lower Akaike Information Criterion (AIC). To test the robustness at different connectivity thresholds, network parameters were computed in a range of 1% to 48% of connectivity densities. Statistical significance was corrected with a false discovery rate (FDR q < 0.05; Benjamini and Hochberg, 1995) throughout the tested range. GAMM models were implemented with the R-package ‘*gamm4*’ (https://CRAN.R-project.org/package=gamm4). In addition, network properties were estimated for each functional network at the 25% connectivity density; significance was corrected with FDR at q < 0.05 for the thirteen functional networks defined in the atlas (Power et al., 2011).

### 2.7 Brain-Cognition inference

A cross-sectional subset of the sample (N = 59; 24 males; age range: 9.67 - 17.98 y.o.) fulfilled a neuropsychological assessment that included a Card Sorting Task (CST; Berg, 1948; Lázaro et al., 2010). The CST measures executive control, reasoning, and learning, but is particularly sensitive to task shifting or cognitive flexibility. Briefly, a set of 64 cards are given one at a time to the participant; each card can be categorized based on color, shape, or the number of figures. The participant has to guess the correct category by putting the card behind four templates, and after that is told if the selection was correct or incorrect. After ten consecutive correct attempts, the category is shifted without informing the participant, thus s/he has to guess again. Several scores are recorded: the score of corrected attempts, perseverations (consecutive errors in the same category), and recurrent perseverations (non-consecutive perseverations in a three attempt span). The total score can be interpreted as a measure of cognitive flexibility; meanwhile, perseverations reflect impulsivity and a lack of executive control and task-switching.

The relationship between the task scores and the functional connectomes was assessed with the Network-Based Statistics approach (Zalesky et al., 2010). This algorithm computes the likelihood of each set of connected edges that surpass an initial statistical threshold (also referred to as clusters of connections, sub-networks, or components). Such likelihood estimation is based on their size (sum of the weights of the edges in this case) and how it compares to a null distribution generated by permutations of the original data. Specifically, the null distribution is the density of the maximum size of the suprathreshold component for each permutation of the original dataset. In this case, the three standardized cognitive scores were fit for each intra- and inter-connection of a connectome of 13×13 functional networks (Power et al. 2011). The model includes the Generalized Additive Model (GAM) spline of the degree at the 25% connectivity density, plus PDS, sex, head-motion, and coil as covariates. A null distribution was computed based on 1,000 permutations setting p < 0.01 for individual connections. GAM splines were implemented with ‘gamm4’ software as well.

## 3. Results

### 3.1 Head-Motion mitigation strategies over functional connectivity

We tested two confounding strategies to minimize the effect of head motion on the brain connectivity estimates; one included the global signal regression (GSR) and the other did not (no-GSR). For the functional connectivity density, both strategies showed a sharp distribution centered on zero. On the other side, graph theory measures showed different associations with head motion after implementing such strategies, but in general for all measures, using GSR resulted in lower associations with head motion, as reflected by their distributions being centered closer to 0 (Figure 2A). Additionally, we tested if these confounding strategies were affected by the Euclidean distance between the nodes for the head motion at the edgewise level, however, both approaches showed a pattern with a slope near to zero (Figure 2B).

**Figure 2.**
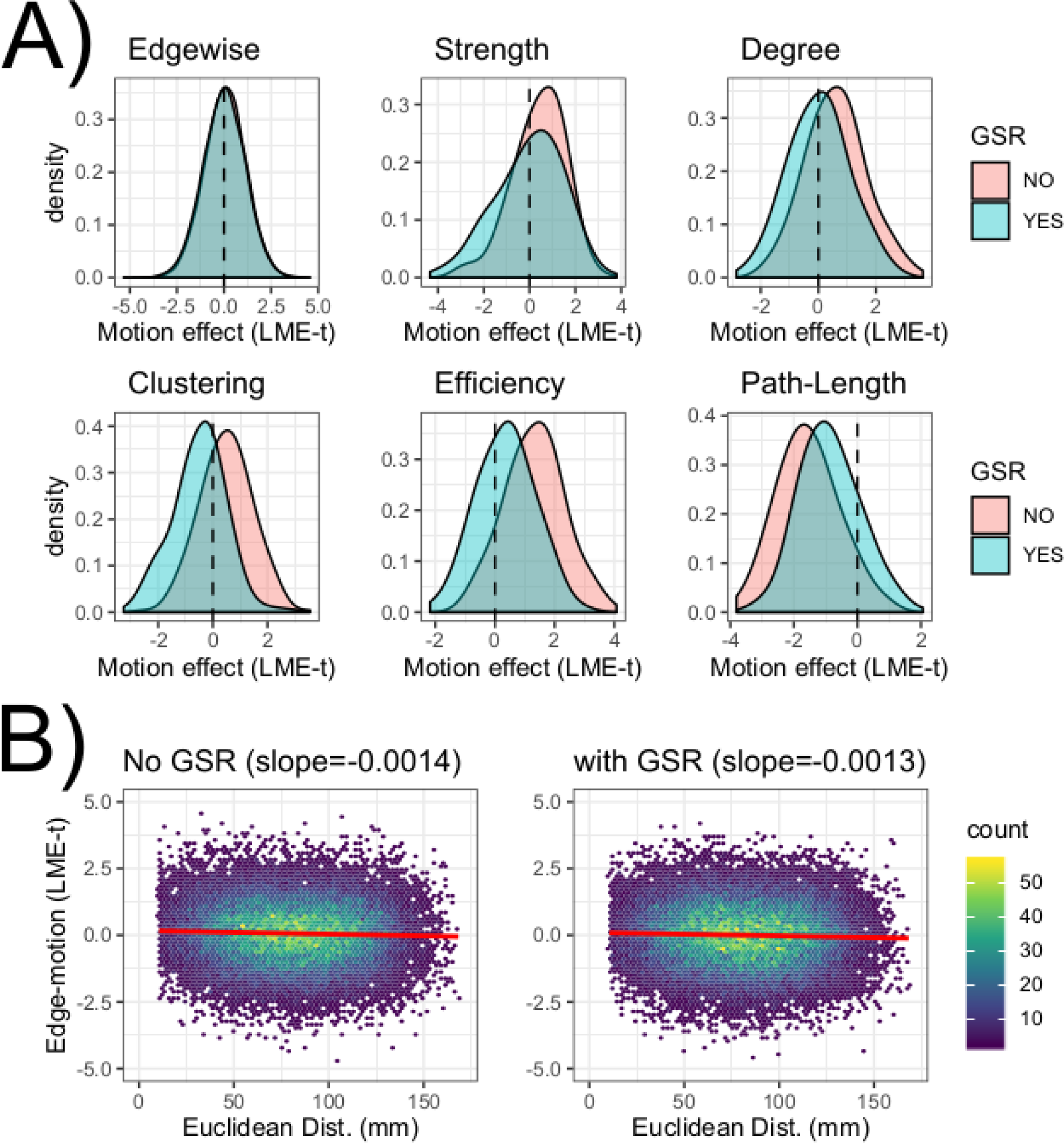
A) Density plots of the linear mixed-effects (LME) t-scores of the relationship between the head motion (measured by the average FD-RMS) and the functional connectivity of edgewise and nodal graph theory variables, for both preprocessing approaches: with and without Global Signal Regression (GSR). B) Hexbin plots of the Euclidean distance between pairs of nodes and the association of head motion with functional connectivity (LME t-scores), for both preprocessing strategies: with and without GSR. Colormap expresses the count of edges with that effect. The fitted line of this relationship is depicted in red.

### 3.2 Neurodevelopmental trajectories

Regarding the pubertal scale along chronological age, males showed a positive linear trend (EDF = 1; F = 674.4; p < 2e-16; GAMM), while females showed a sigmoid shape with faster growth than males around 12 years of age (EDF = 2.96; F = 456; p < 2e-16; GAMM) (Figure 3).

**Figure 3.**
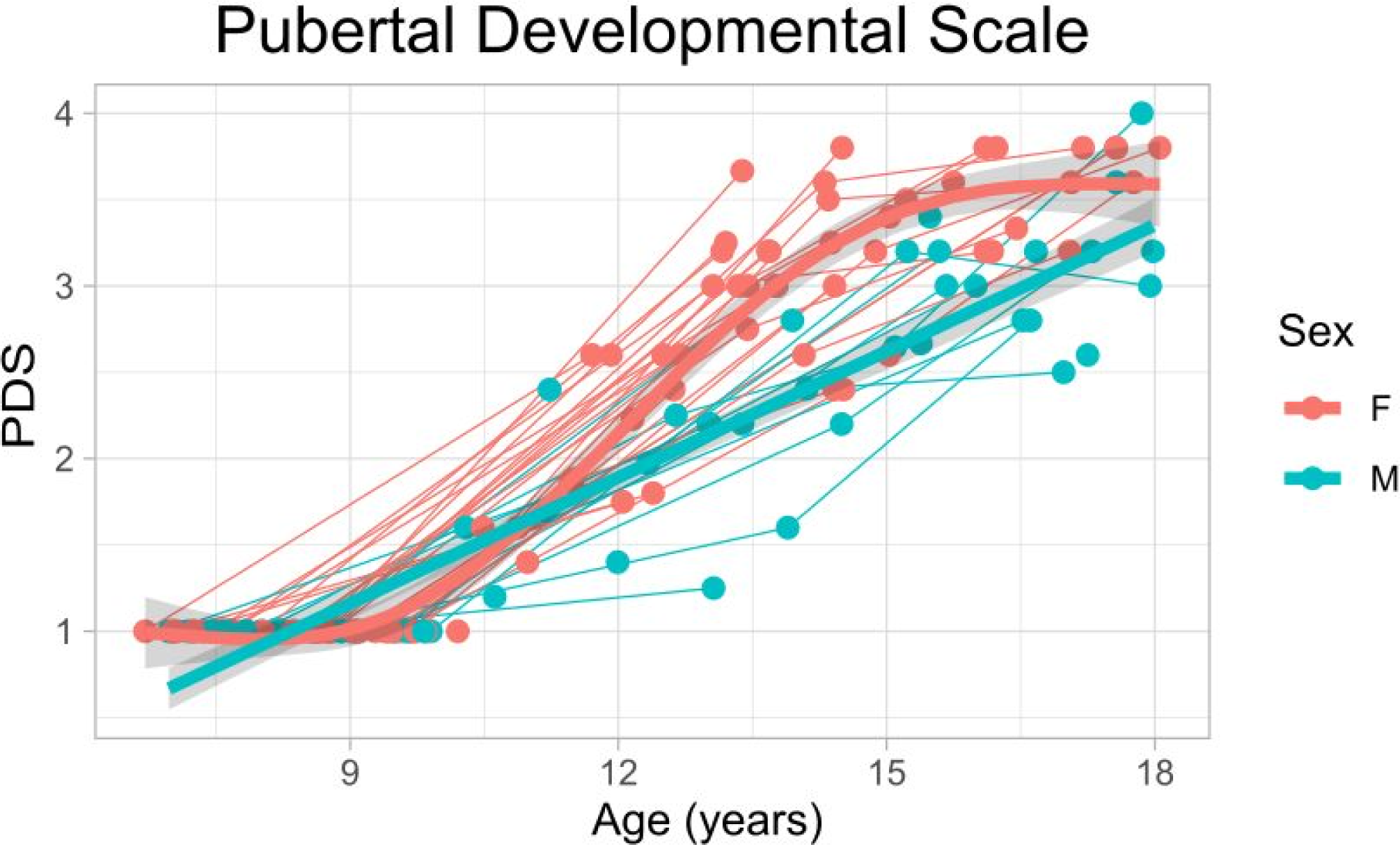
Pubertal Development Scale scores (PDS) along age. Thin lines represent individual trajectories; thick lines represent GAMM splines per sex group (with 95% confidence-interval shadow). Abbreviations: female (F), male (M).

When comparing the adjustment of the four different GAMM models (age, PDS, age-by-sex, PDS-by-sex) for each network measure at different connectivity densities, we found that for every measure, the models including PDS showed the best fit for the majority of connectivity densities (Figure 4). In addition, in most of these cases, the GAMM spline of the PDS is significant after the correction for multiple comparisons (FDR q < 0.05).

**Figure 4.**
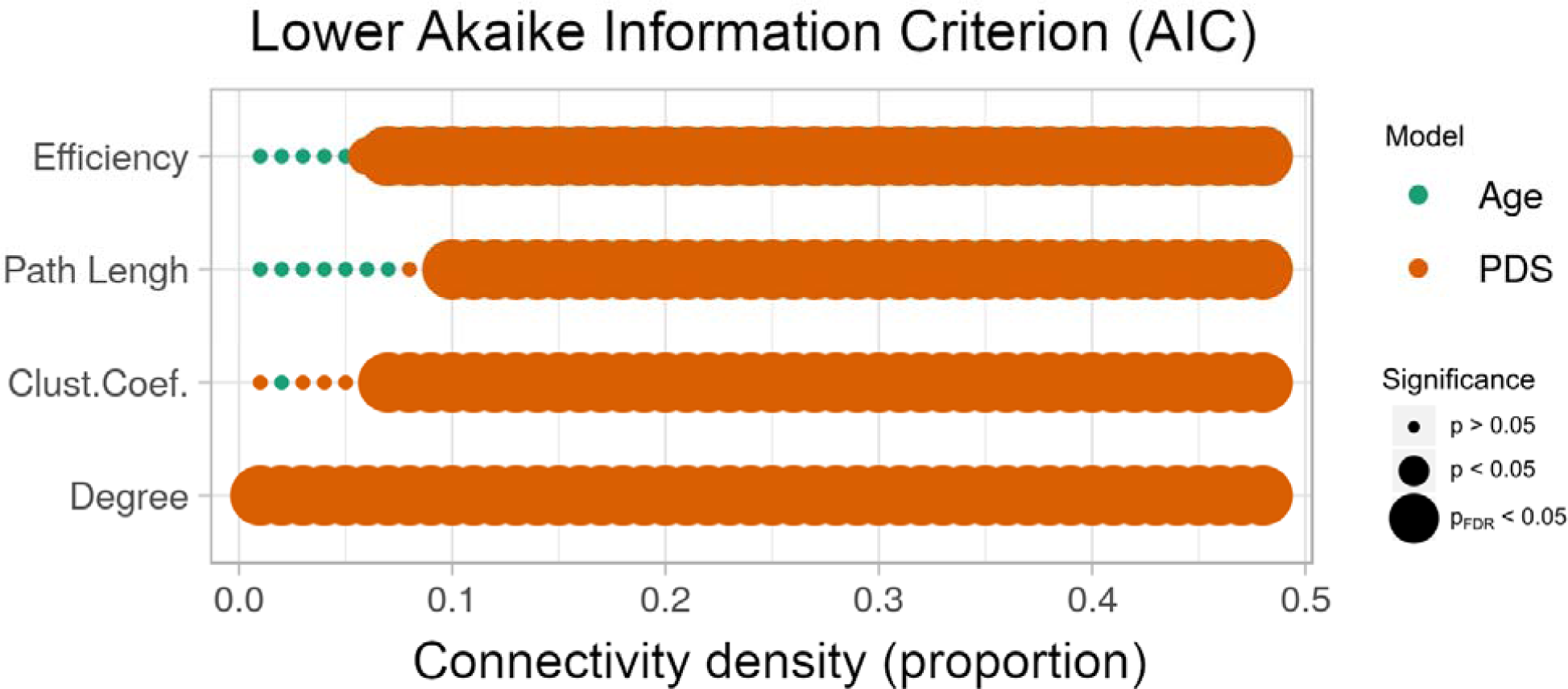
The model (age, PDS, age-by-sex, PDS-by-sex) with lower Akaike Information Criterion (AIC) is depicted for each graph theory measure (y-axis) and every connectivity density (x-axis) here explored. The significance level is depicted with circle size.

Taking a representative connectivity density of 25%, the PDS fits better than the rest models (Table 1), which is consistent with the majority of other densities (Figure 4). At this threshold, we can observe the neurodevelopmental trajectories via GAMM splines, for the four network measures the trend along the PDS was quadratic, with a point of inflection around the level 2 of the pubertal scale. In particular, the degree, clustering coefficient, and efficiency showed a convex pattern, where these measures increase in value after the turning point (Figure 5), while the path-length showed the inverse pattern, a concave curve (Figure 5).

**Table 1.**
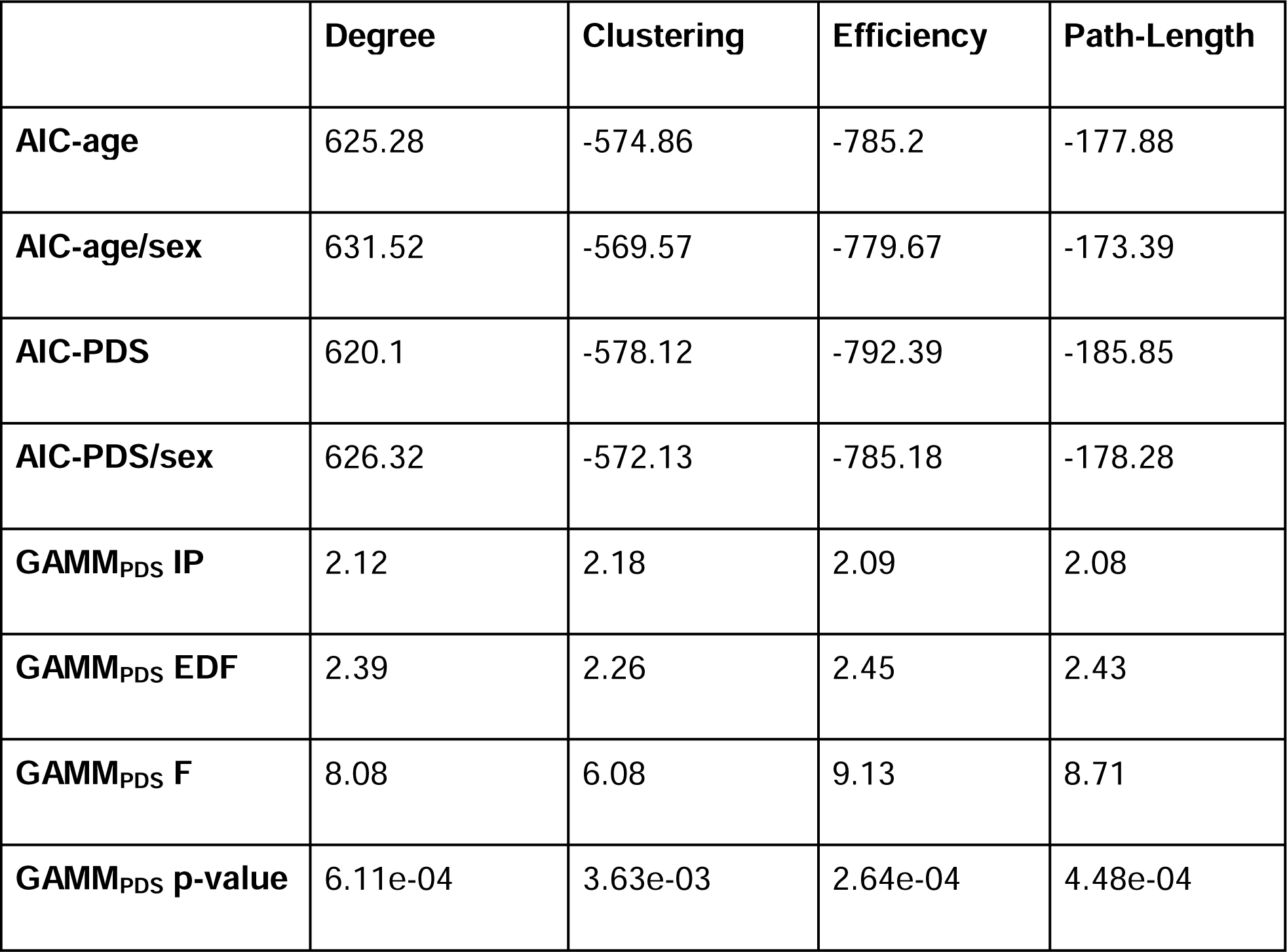
For every graph theory measure computed at the 25% connectivity density, the Akaike Information Criterion (AIC) for every graph theory measure of the four Generalized Additive Mixed Models (GAMM) models tested: age, age-by-sex, Pubertal Development Scale (PDS), and PDS-by-sex; plus the PDS GAMM’s spline inflection point (IP), estimated degrees of freedom (EDF), F-value, and p-value.

**Figure 5.**
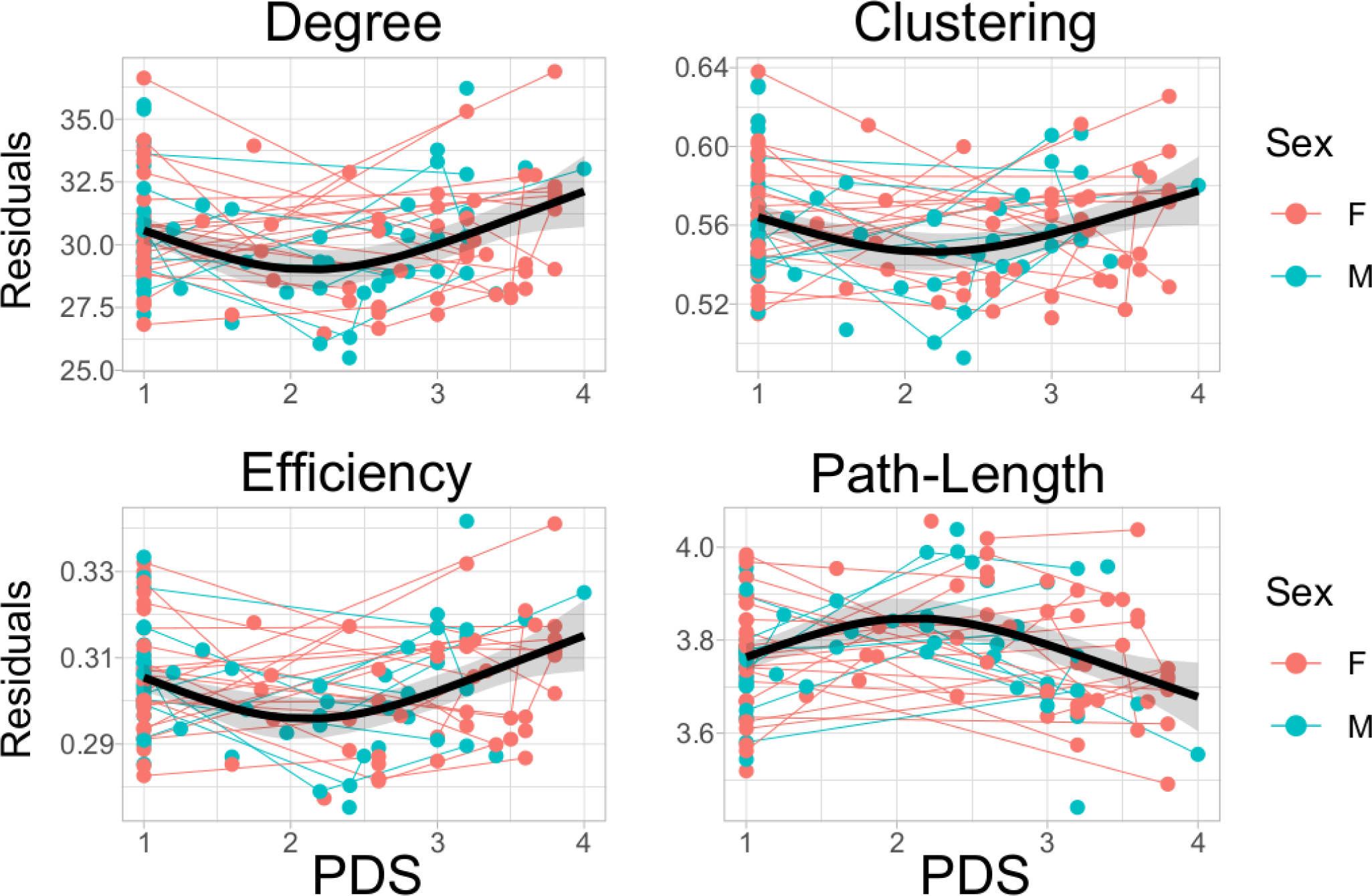
Scatter-plots of the best model at the 25% connectivity density for each graph theory residuals (after regressing out in-scanner motion and head-coil). Degree (top-left), clustering coefficient (top-right), efficiency (bottom-left), and characteristic path length (bottom-right) in relation to the pubertal scale (PDS). Thin lines represent individual trajectories; thick black lines represent the sample GAMM splines (with 95% confidence-interval shadow).

At the functional network level, neurodevelopmental effects along the pubertal stage showed the same quadratic pattern found in the whole-brain analysis, but mainly in frontal and parietal areas for every graph theory measure. Particularly, for the degree, stronger effects were found in the default mode (DMN), fronto-parietal (FPN), and ventral attention (VAN) networks (Table 2), with a positive effect after the point of inflection of the curve (Figure 6). The clustering coefficient showed a similar pattern, but with no significant effects after correcting for multiple comparisons. For efficiency, all the functional networks that involve frontal and parietal areas, plus the cerebellar (CBL), subcortical (SUB), and visual (VIS) networks showed a strong effect along the PDS (Table 2). While for the path-length, as expected, the effects were inverse to those of efficiency, with a decrease in magnitude after the point of inflection (Table 2; Figure 6).

**Table 2.**
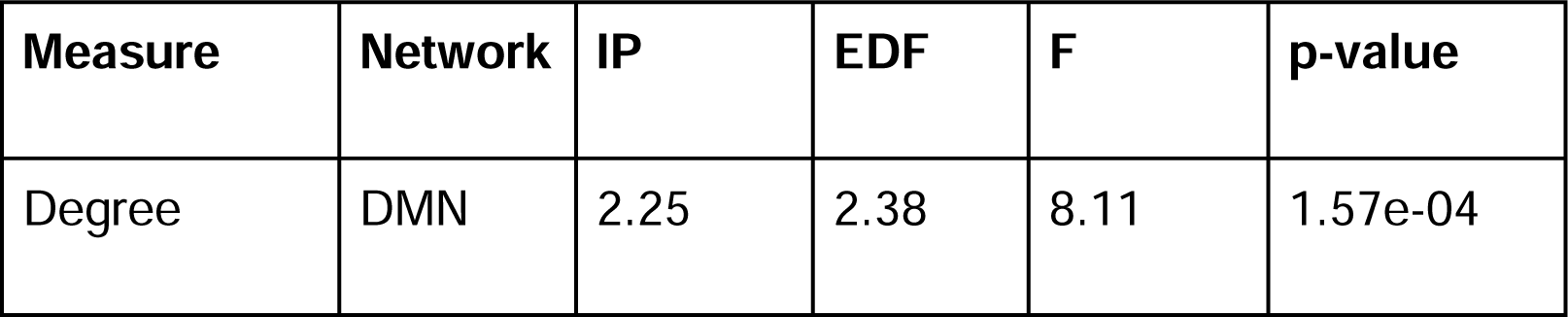
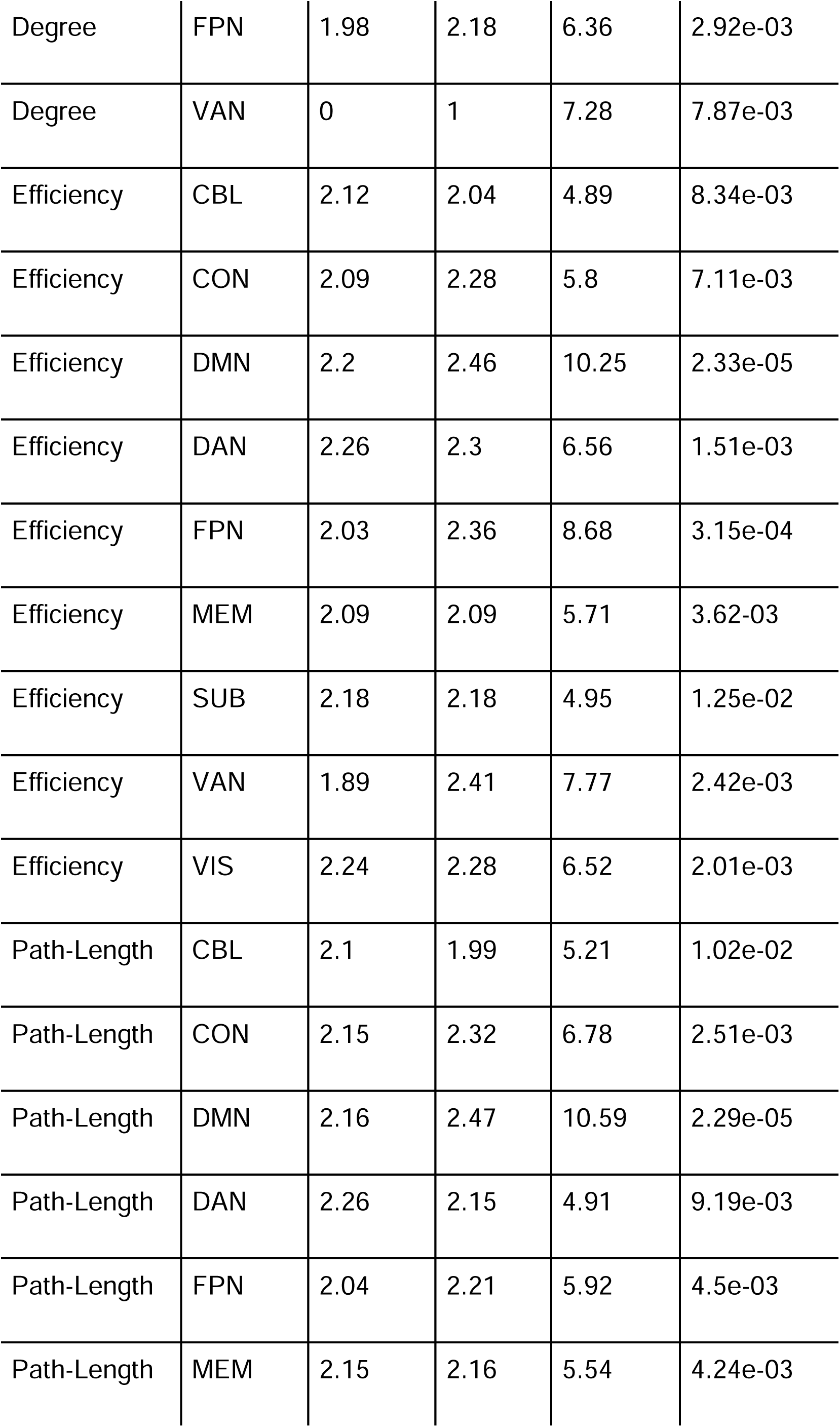
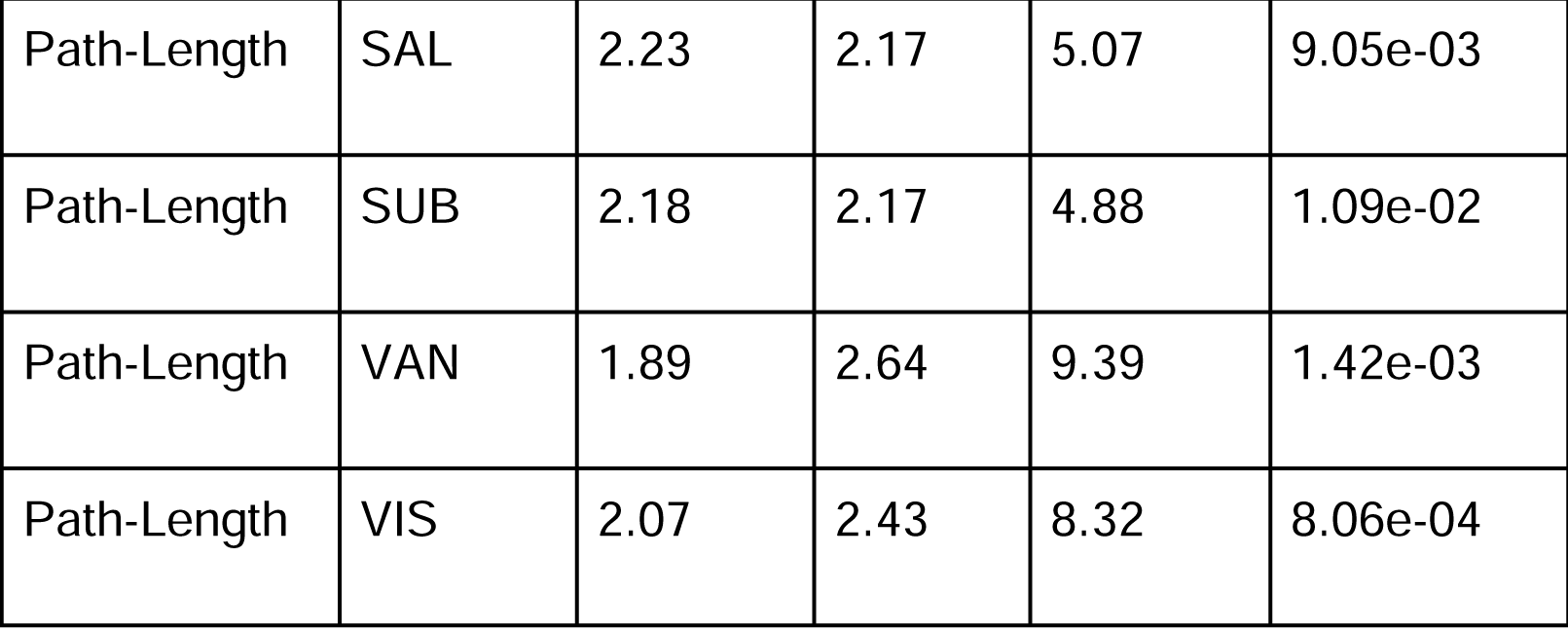
Graph theory measures with significant splines for PDS at the functional network, corrected for FDR q < 0.05. Values for PDS GAMM’s spline inflection point (IP), estimated degrees of freedom (EDF), F-value, and p-value. Network abbreviations: cerebellar (CBL), cingulo-opercular (CON), dorsal attention (DAN), default-mode (DMN), fronto-parietal (FPN), memory (MEM), salience (SAL), subcortical (SUB), ventral attention (VAN), visual (VIS).

| Measure | Network | IP | EDF | F | p-value |
| --- | --- | --- | --- | --- | --- |
| Degree | DMN | 2.25 | 2.38 | 8.11 | 1.57e-04 |
| Degree | FPN | 1.98 | 2.18 | 6.36 | 2.92e-03 |
| Degree | VAN | 0 | 1 | 7.28 | 7.87e-03 |
| Efficiency | CBL | 2.12 | 2.04 | 4.89 | 8.34e-03 |
| Efficiency | CON | 2.09 | 2.28 | 5.8 | 7.11e-03 |
| Efficiency | DMN | 2.2 | 2.46 | 10.25 | 2.33e-05 |
| Efficiency | DAN | 2.26 | 2.3 | 6.56 | 1.51e-03 |
| Efficiency | FPN | 2.03 | 2.36 | 8.68 | 3.15e-04 |
| Efficiency | MEM | 2.09 | 2.09 | 5.71 | 3.62e-03 |
| Efficiency | SUB | 2.18 | 2.18 | 4.95 | 1.25e-02 |
| Efficiency | VAN | 1.89 | 2.41 | 7.77 | 2.42e-03 |
| Efficiency | VIS | 2.24 | 2.28 | 6.52 | 2.01e-03 |
| Path-Length | CBL | 2.1 | 1.99 | 5.21 | 1.02e-02 |
| Path-Length | CON | 2.15 | 2.32 | 6.78 | 2.51e-03 |
| Path-Length | DMN | 2.16 | 2.47 | 10.59 | 2.29e-05 |
| Path-Length | DAN | 2.26 | 2.15 | 4.91 | 9.19e-03 |
| Path-Length | FPN | 2.04 | 2.21 | 5.92 | 4.5e-03 |
| Path-Length | MEM | 2.15 | 2.16 | 5.54 | 4.24e-03 |
| Path-Length | SAL | 2.23 | 2.17 | 5.07 | 9.05e-03 |
| Path-Length | SUB | 2.18 | 2.17 | 4.88 | 1.09e-02 |
| Path-Length | VAN | 1.89 | 2.64 | 9.39 | 1.42e-03 |
| Path-Length | VIS | 2.07 | 2.43 | 8.32 | 8.06e-04 |

**Figure 6.**
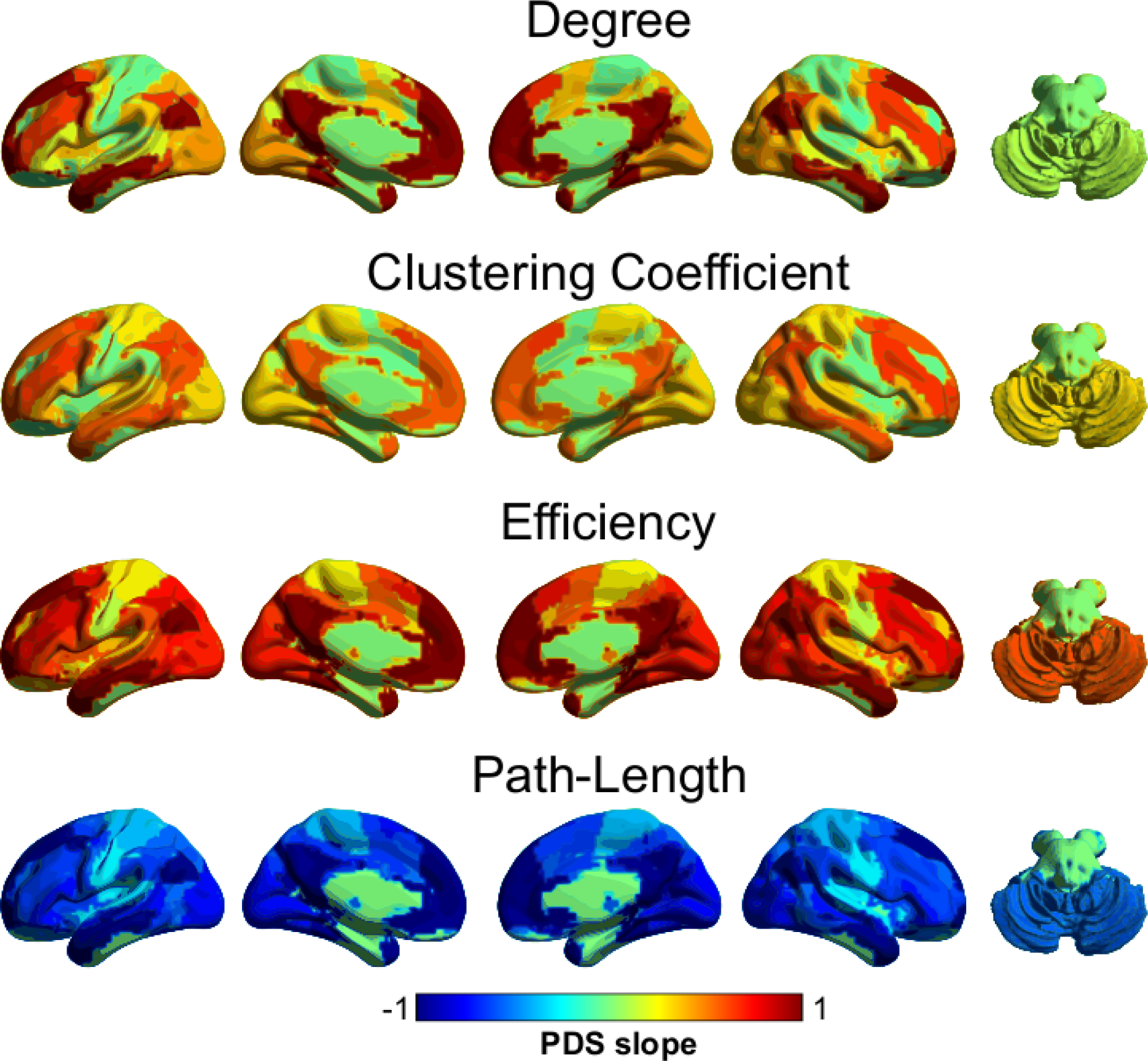
Brain map (Xia et al., 2013) of the GAMM spline slope after the point of inflection for each graph theory measure. Mapping was based on ROI corresponding consensus area according to Power et al. (2011).

### 3.3 Brain-cognition relationships

Regarding the relationship of the functional connectome and cognitive performance, NBS identified two significant sets of network interactions that relate with performance in the CST assessment (Figure 7). Specifically, higher connectivity between the cingulo-opercular network (CON), dorsal attention (DAN), default mode (DMN), and fronto-parietal (FPN) networks was associated with higher scores in the CST (p_FWE_ = 0.0458; Figure 7A). In addition, lower number of recurrent perseverations were associated with higher connectivity of the cingulo-opercular network (CON), within itself and with the memory retrieval network (MEM); in contrast, higher number of recurrent perseverations was associated with higher connectivity of the sensorimotor-hand network (SMH) (p_FWE_ = 0.0236; Figure 7B).

**Figure 7.**
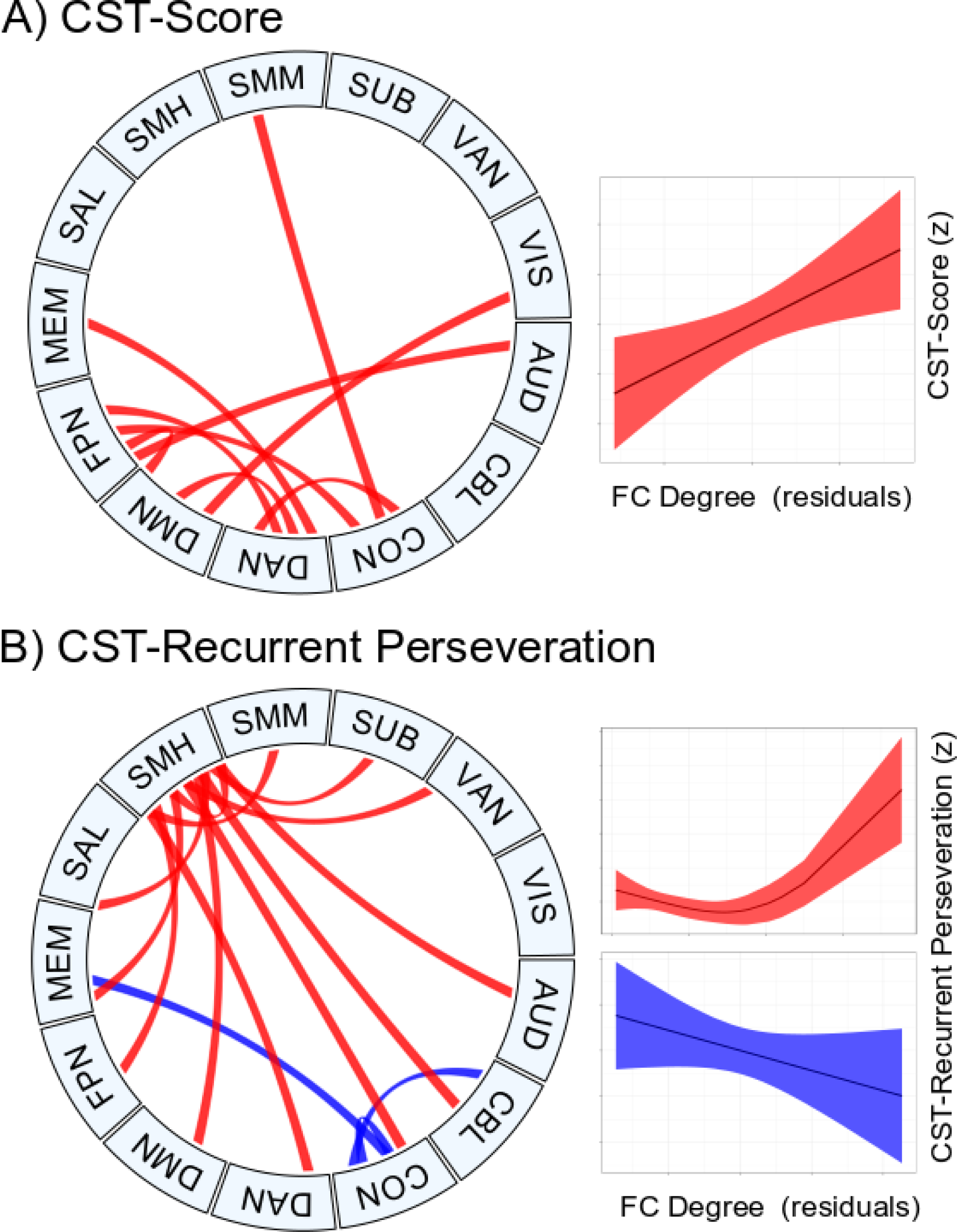
Chord diagrams (Z. Gu et al., 2014) of the GAM Network-Based Statistics of the relationship between the functional connectivity degree and the score (A) and the recurrent perseveration (B) of the Card Sorting Task (CST). These components were significant after correction for multiple comparisons with a p_FWE_ of 0.0458 and 0.0236, respectively.

## 4. Discussion

By exploring a longitudinal sample and with a strict control of the effects of head motion, the present study showed that the brain functional organization along adolescence follows non-linear puberty-dependent trajectories, with an inflection point around the beginning of puberty, when the first signs of body changes appear (level 2 of the PDS). The frontal and parietal brain regions exhibited the most dramatic rates of change after the inflection point. Finally, a specific pattern of network interactions reflected higher performance in a cognitive task associated with executive functions, mainly cognitive flexibility. Such a pattern includes the higher integration of fronto-parietal networks and the functional segregation of primary networks.

### 4.1 Head-motion effects on graph theory

Taking two rigorous confounding strategies to mitigate the effect of head motion on the network estimates, considering or not considering the global signal regression (GSR), we found that both strategies properly account for this effect for the edgewise functional connectivity, even when considering the Euclidean distance between nodes. When exploring such associations for the graph theory measures, they are closer to zero for the approach using GSR (Figure 2). In addition to the confounding strategies at the subject level, the average head motion for each subject was included at the group level analyses, to further minimize its effect on the identified trajectories.

Previous pediatric studies have found similar results of the motion effects on edgewise functional connectivity, with the 36 parameters and the *aCompCor* being two of the most effective strategies amid other widely used approaches (Ciric et al., 2017; Taymourtash et. al, 2019). However, when considering graph theory measures only one previous work in adult populations has tested the in-scanner motion effects. Aurich et al. (2015) tested several preprocessing strategies of motion mitigation on graph theory measures, finding that censoring of “spikes” reduces the association with the head-motion estimates, even lower than GSR. Nevertheless, they did not test more rigorous strategies such as derivative, quadratic trends, PCA, etc. and the inclusion of several mitigators in the same strategy, such as we did in our study. Therefore, when considering a rigorous amount of motion regressors the head motion effect on graph theory measures is effectively controlled for the degree, clustering, and efficiency, but less for path-length (Figure 2A).

### 4.2 Puberty and neurodevelopmental trajectories

Pubertal status showed a faster growth rate in females compared to males. Although self-reported questionnaires are sensitive to inaccuracies (Dorn et al., 2006), the pattern observed has been replicated in other populations (Braams et al., 2015; Wierenga et al., 2018; van Duijvenvoorde et al., 2019). Pubertal scale scores showed a faster growth rate in females around level 2 and 12 years of age compared to males. Also the PDS fits better to brain functional organization measures than chronological age, with inflection points at the first signs of puberty (level 2 of the PDS). Functional centrality, segregation, and efficiency increases after the inflection, while path-length decreases after such a point, being more prominent in frontal and parietal functional networks in all cases.

Notably, turning points coincide with the start of puberty. No previous work has focused on the functional organization of the brain network; however, animal studies have demonstrated how the start of the hormone events of puberty provoke plastic changes in the structure and function of the brain (Laube et al., 2020). Pubertal hormones at the onset of puberty are related to an increase of the synaptic pruning in frontal areas (Drzewiecki et al., 2016), as well as a growth in inhibitory neurotransmission in these areas while not in sensory regions (Piekarski et al., 2017). These results evince the changes in the brain connectivity patterns at lower scales following the onset of puberty, particularly in association areas. In addition, human studies have shown the relevance of pubertal status in gray matter (Goddings et al., 2014), white matter microstructure (Genc et al., 2017), and even fronto-subcortical functional connectivity (van Duijvenvoorde et al., 2019). Other neuroimaging studies have found lower gray matter density in cortical and subcortical areas in adolescence with higher pubertal status when controlling for chronological age (Peper et al., 2009; Bramen et al., 2011). Similarly, puberty has been related with higher white matter density (Perrin et al., 2009; Herting et al., 2012), which can be interpreted as a maturity advance (Gogtay et al., 2004; Lenroot et al., 2007; Lebel and Beaulieu, 2011; Coupé et al., 2017). Also, the same effects of lower gray matter and greater white matter density have been related to blood levels of testosterone (Paus et al., 2010). These results, along with those presented here, demonstrate how the onset of puberty greatly impacts brain structure and function. In particular, we showed an increase in the efficiency of the brain network. Furthermore, the pubertal markers have implicit sex dimorphisms that explain the functional neurodevelopment better than chronological age or sex-age effects. Hence, it is highly recommendable that developmental studies in adolescence include estimations of the pubertal stage.

Regarding the whole-brain functional network, the results showed an increase after the onset of the puberty in brain functional centrality, segregation, efficiency, and integration (i.e., the inverse of path length). Although no studies have tested the connectome-puberty relationship, previous cross-sectional studies have shown a widespread linear increase in functional connectivity along with chronological age during adolescence (Hwang et al., 2012; Satterthwaite et al., 2013a; Marek et al., 2015). However, these studies differ in how this increase impacts functional segregation or integration. Satterthwaite et al. (2013a) emphasized that the linear increase in functional connectivity along with age is more prominent within functional networks, which implies an increase in functional segregation. On the other hand, Marek et al. (2015) showed a decline in the functional connectivity from childhood to adolescence followed by an increase in late adolescence, particularly between network connectivity, i.e., higher functional integration. The latter fits with our nonlinear results for degree and path length; however, they did not show the same pattern regarding functional segregation. In summary, previous studies differ in the evolution of the connectomic functional segregation and integration, potentially due to differences in the methodological approaches; nevertheless, the implementation of nonlinear splines in the present study was able to extract the main effects previously reported. Therefore, our results contribute to converging those earlier findings, and also to emphasize the relevance of taking nonlinear approaches in developmental contexts.

Concerning the development of specific functional networks, significant effects were found in the degree, efficiency, and path length after controlling for multiple testing, mostly involving attention and task control related systems: dorsal attention (DAN), default mode (DMN), cingulo-opercular (CON), fronto-parietal (FPN), memory retrieval (MEM), salience (SAL), and ventral attention (VAN). In particular, those effects were more prominent than those found in the whole-brain inferences. These results are congruent with previous work, which showed an increase in within and between task control networks. Hwang et al. (2012) reported higher functional connectivity in adolescence within frontal and between frontal and parietal areas, while Marek et al. (2015) found an increase in functional integration in the same task control networks along this period. Our results combined with previous studies coincide with the strengthening of cognitive control brain networks, which strengthens along with the control-related behavior (Mennes et al., 2010; Baum et al., 2017), but ours highlight the relevance of puberty onset for the turning point of brain functional organization, particularly those that include frontal and parietal regions.

### 4.3 Brain-cognition effects

Even controlling for the pubertal effects, higher connectivity between attention and task control networks was associated with higher scores and lower error rates in a task evaluating cognitive flexibility. In contrast, this pattern was inverse when considering the sensorimotor-hand network (SMH). Previously, Marek et al. (2015) demonstrated that higher functional integration (participation coefficient) between the cingulo-opercular and salience networks was associated with better accuracy on inhibition tasks. This shows that the integration of attention/task-control compared to primary/unimodal systems not only reflects the brain organization development, but also that such patterns are very relevant for cognitive control and cognitive flexibility.

## 5. Limitations

A potential limitation in this study was the missing values in the pubertal scale scores. Such information was not collected in the first sampling, which included participants with ages lower than 10 years old. These values were set to score 1, as similar values have been shown in comparable samples (Hibberd et al., 2015; van Duijvenvoorde et al., 2019). In addition, a minor number of values (n=8) in older participants were estimated via a GAMM approach. However, the adjustment of the GAMM splines to the observed data (R2-adj = 93.6) together with the reproducibility of similar spline shapes in other samples, lead us to consider the estimated PDS scores as a reliable measure of pubertal status.

Another limitation is the absence of assessment of the menstrual cycle in females. Although the menstrual cycle has not been associated with changes in functional connectivity in fronto-parietal networks (Hjelmervik et al., 2014), menstrual cycle phases are related to structural and functional changes regarding subcortical structures (Lisofsky et al., 2015; Hidalgo-Lopez et al., 2020).

## 6. Conclusion

The present work shows that the brain functional organization follows a nonlinear trend that fits the pubertal status during the adolescent period. In particular, brain functional centrality, segregation, efficiency, and integration increased after the onset of the pubertal markers, this effect being more pronounced in regions related to attention and task control networks. In addition, higher connectivity between those regions was associated with better cognitive flexibility performance. In summary, pubertal status along with longitudinal and nonlinear patterns contribute to better description of the developmental trajectories of the functional connectome, even converging previous discrepancies regarding both segregation and integration increase in adolescence.

## Acknowledgements

We are extremely grateful to the children and parents that voluntarily participated in this study. We are grateful to Nuri Aranda López, Leonor Casanova Rico, Deisy Gasca Martínez, Leopoldo González-Santos, Ma. de Lourdes Lara Ayala, Juan J. Ortiz and Erick H. Pasaye for their technical support. We are also grateful to Roberto E. Mercadillo for his support in the psychological assessment and Michael C. Jeziorski for editing the manuscript. The authors thankfully acknowledge the imaging resources and support provided by the “Laboratorio Nacional de Imagenología por Resonancia Magnética”, CONACYT network of national laboratories. Zeus Gracia Tabuenca is a doctoral student of “Programa de Doctorado en Ciencias Biomédicas, Universidad Nacional Autónoma de México” (UNAM) and received a fellowship (330142) from “Consejo Nacional de Ciencia y Tecnología” (CONACYT). CONACYT had no role in study design, data collection, analyses nor writing the manuscript. This research was partially supported by grant UNAM-DGAPA-PAPIIT IN212219 to SA.

